# Rhizobia independently adapt to soil and legume host environments, but soil conditions influence the abundance of high-quality partners

**DOI:** 10.64898/2025.12.08.692960

**Authors:** Alejandra Gil-Polo, Regina B. Bledsoe, McCall Calvert, Lillian Cherry, Brendan Epstein, Rebecca Fudge, Jennifer Harris, Peter Tiffin, Liana T. Burghardt

**Affiliations:** The Pennsylvania State University, University Park, USA; University of Pennsylvania, Philadelphia, USA; The Nature Conservancy, Arlington, Virginia, USA; University of Minnesota, USA

**Author notes:** Author for correspondence:* Liana T. Burghardt, Department of Plant Science, 109 Tyson Building, University Park, PA 16802, 1-(.

**Keywords:** Mutualism, GWAS, *Medicago truncatula*, *Sinorhizobium meliloti*, bacterial fitness correlations, soil, temperature, cold, salinity, heat, symbiosis, free-living

## Abstract

Rhizobia live as free-living microorganisms in the soil and in association with legume hosts. Both environments exert selective pressures on rhizobia, influencing the reproductive success of individual strains (e.g., fitness). The soil, a heterogeneous and fluctuating environment, is often overlooked, and little is known about whether selection in the soil influences the outcomes of the rhizobia-legume mutualism. We exposed a mixture of 68 *Sinorhizobium meliloti* strains to eight soil treatments (temperature, osmotic, and texture perturbations) and selection by two *Medicago* plant hosts. We found that cold (4°C) and warm (32°C) temperatures, as well as salt addition, had the strongest effects on diversity, community composition, or population size. Strain relative fitness was strongly positively correlated among soil treatments, except for cold. Genome-wide association analysis revealed a complex genetic architecture for soil fitness. In contrast, when comparing rhizobial fitness between soil and host environments, we found minimal strain fitness correlations, suggesting independent genetic bases and habitat-specific adaptations. Lastly, by examining the relationship between rhizobial fitness in the soil and their nitrogen-fixing plant benefits, we found that soil selection influenced the relative abundance of high- and low-quality strains; However, whether these effects were positive or negative for the plant was host dependent. Our results suggest that rhizobial evolution in soil and host are largely independent, but soil selection can alter mutualism benefits.

**IMPORTANCE:** Rhizobia-legume mutualism is crucial for introducing nitrogen into agricultural and natural ecosystems, and rhizobia persistence in the soil is an important component of agroecosystems. However, we know little about how individual strains of rhizobia persist and adapt to this environment, especially in the context of the soil’s spatial and temporal variations (temperature, moisture, and soil texture). We found that rhizobia similarly adapt to abiotic soil conditions but their reproductive success in the soil is independent from their reproductive success in the host. Intriguingly, we found that certain soil conditions increase (or decrease) the relative abundance of more effective nitrogen-fixing strains. Understanding how rhizobia adapt to diverse environments is crucial for developing effective bioinoculants that maintain high persistence in the soil while are also highly competitive to colonize the host and are beneficial to the plant.

## INTRODUCTION

Facultative mutualists can survive and reproduce with or without their mutualistic partner (1). This flexibility means that each partner faces selective pressures during both their free-living and partner-associated phases. A well-studied example of facultative mutualism is the interaction between rhizobia bacteria and leguminous plants. Rhizobia alternate between existing as free-living microbes in the soil, where they live as saprophytes, and as mutualistic symbionts within the root nodules of plants, where they fix atmospheric nitrogen in exchange for sugars derived from photosynthesis (2–4). However, because much of the research on this mutualism has focused on understanding microbial interactions with host plants (5), our understanding of the factors affecting rhizobial fitness in the soil and how selection in this habitat could cascade to affect the functioning of the mutualism remains limited (6, 7).

The soil is a crucial and variable habitat for rhizobia. Rhizobia can spend many years living in the soil, either because their host is not present (3, 8–12) or because of the low probability, estimated at one in a million, that they would form a nodule even when their host is present (3). In multistrain communities, natural isolates of the same rhizobial species display varied fitness in the soil, meaning that strains differ in their survival and reproduction (6, 13). However, these studies have primarily been conducted at room temperature, and don’t account for the substantial variability in soil, which is spatially structured (e.g., texture and moisture) (14) and varies across seasons (15). Research that has considered factors such as temperature, salinity, drought, and soil texture has demonstrated their impact on the size of rhizobial populations, but these studies have typically focused on single strains (16–19) or have estimated rhizobia numbers at the broad genus level in field assessments (20, 21). This study investigates how soil properties function as selective agents for a diverse rhizobial population, the consistency of selection across soil perturbations, and the genetic basis of rhizobial fitness in the soil.

Nodules offer rhizobia the potential for exponential growth (22) and accumulating resources that increase survival (23); therefore, rhizobial strains compete to associate with the host (24–26). Consequently, rhizobial fitness in hosts reflects the cumulative effect of the selection in the soil, rhizosphere, root surface, infection thread, and after they have established inside the nodules (2). It remains unknown whether rhizobial fitness across these environments covaries, i.e., whether strain fitness in the soil and in the host’s nodules is correlated (27). A positive correlation in fitness between the soil and host environments would enhance adaptation to both settings, while a negative correlation may limit adaptation (28, 29). Negative correlations could also explain the considerable genetic variation in rhizobial partner quality observed within many populations (7). To address this gap, we utilize trait correlations and Genome-Wide Association (GWA) analyses to assess whether fitness traits covary between habitats and whether this correlation is due to a shared genetic basis (30, 31).

In this paper, we assess whether adaptations of rhizobia to various abiotic perturbations in the soil correlate across different soil conditions and with two traits related to mutualism: rhizobial fitness in the host and rhizobial partner quality. We used 68 strains of *Sinorhizobium meliloti* whose genetic diversity served as the raw material for selection. The genomes of these strains have been sequenced, which allowed us to use genomic markers to discriminate among them using a Select and Resequence approach and to identify genomic regions associated with microbial fitness in the soil and hosts using Genome-Wide Association Studies (GWAS) (32, 33). We assessed the fitness of these strains in eight soil mesocosm environments (Fig. 1), using two *Medicago* accessions, *M. truncatula* (A17) and *M. littoralis* (R108), which select for different *Sinorhizobium meliloti* strains (6). Specifically, we ask: 1) How do soil abiotic factors affect rhizobial population size, community composition, and diversity? 2) To what extent do individual strains show consistent fitness patterns across soil conditions and between soil and host environments? and 3) How is rhizobial fitness during the free-living soil stage related to the benefits they provide to their hosts?

**Figure 1:**
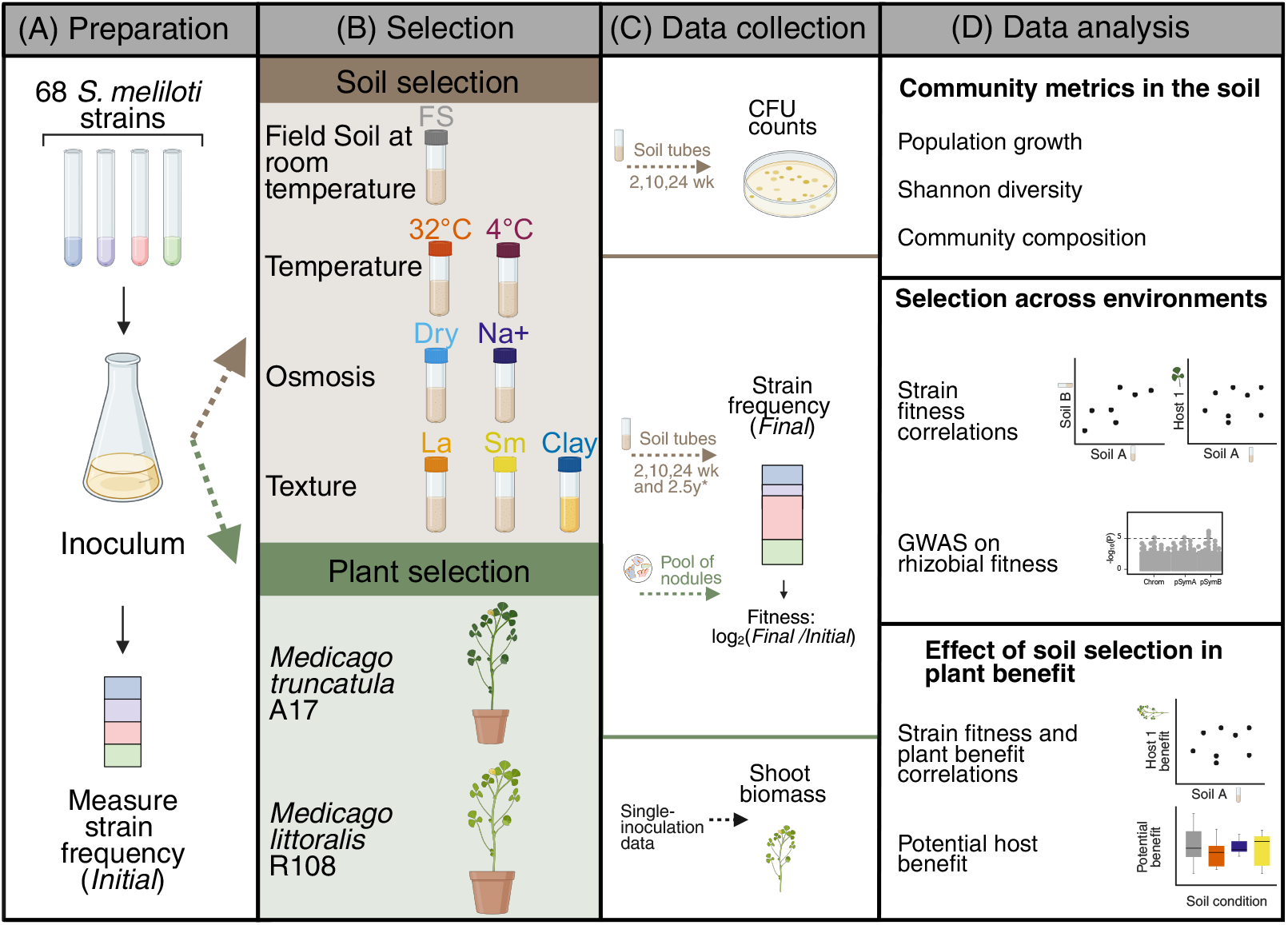
Design of the soil and plant inoculation experiments. (A) We prepared a mixed inoculum using 68 strains of *Sinorhizobium meliloti*. (B) We exposed the mixture to soil and plant selection. For the soil selection, we inoculated soil tubes, imposed seven abiotic treatments, and sampled five replicate tubes at three timepoints (2 weeks, 10 weeks, and 24 weeks). For plant selection, we inoculated six replicate pots of each of two host genotypes, *Medicago truncatula* (A17) and *Medicago littoralis* (R108). To ensure nodule pool sizes of more than 100 nodules, each pot had ~ 10 plants. (C) From the soil samples, we collected the counts of colony-forming units (CFU) using dilution plating. From both soil samples and a pool of nodules, we measured the relative frequency of rhizobial strains and estimated their fitness. From previously published data (6), we obtained the dry shoot biomass of A17 and R108 inoculated with the single strains used in this paper. (D) First, we asked whether abiotic perturbations in the soil affect the following community metrics: population growth, Shannon diversity, and the community composition. Second, we assessed the strain fitness correlations across environments and identified candidate variants for strain fitness using a genome-wide association study (GWAS). Lastly, we asked whether soil tended to enrich for high- or low-quality partners based on the shoot biomass from single inoculation data. For this evaluation, we used two strategies: calculating the correlations between strain fitness in the soil and the plant benefit, or calculating the potential host benefit. FS-control or FS (Field Soil at room temperature), 32°C (Warm treatment at 32°C), 4°C (Cold treatment at 4°C), Dry (Drought treatment), Na+ (High salinity treatment), La (soil with large particle size), Sm (soil with small particle size), Clay (Clay treatment). * For the FS-control treatment, we also sampled the 2.5-year time point. Created in BioRender. Burghardt, L. (2025) https://BioRender.com/qk4b7hi

## MATERIALS AND METHODS

### Experimental design and data collection

We created a mixed inoculation of 68 *Sinorhizobium meliloti* strains selected to capture a majority of genetic diversity found among a sample of 153 strains sequenced by Epstein et al.,(2018) (34). Each strain differed from the others by a minimum of 1000 SNPs. To create an inoculant with approximately equivalent amounts of each strain, we grew each strain in 3mL of Tryptone yeast media (6 g tryptone, 3 g yeast extract, 0.38 g CaCl_2_ per L) at 28°C and 200 rpm for 3 days. We then combined the 68 cultures to form the initial inoculum. Four 1 mL aliquots were pelleted to assess strain frequency in the initial community (Fig. 1A).

To create the soil mesocosm, we added 10 grams of field soil to 25 × 100 mm test tubes. All treatments used sand/loam field soil collected from an agricultural field on the University of Minnesota campus, except for the Clay treatment, which utilized a field soil with a higher clay content. We autoclaved the tubes twice using a 30-minute liquid cycle, adding 3 mL of H_2_O each time to saturate the soil during autoclaving (this 3 mL evaporated each time). To each tube containing 10 g of field soil, we then added 10 μL of the mixed inoculum diluted in 1.99 mL of a 0.85% w/v NaCl solution (~10^7^ rhizobia). To maintain the moisture level, we weighed each tube each week. When a tube lost ~ 1.5 g (12.5% of the total weight and ~50% of the water weight), we added 1.5 mL of sterile H_2_O to bring them back to their original weight, except for the drought treatment (Dry), which we allowed to lose 3 g (25% of the total weight and ~100% of the water weight) before adding 1.5 mL sterile water. We maintained all tubes at room temperature (~22°C), except for the cold treatment (4°C), which was kept in a refrigerator set to 4°C, and the warm treatment (32°C), which was kept in an incubator set to 32°C. For the salinity treatment (Na+), we replaced 1 mL of the H_2_O with 300 mM NaCl at the onset of the experiment. To establish the texture treatments, we sieved the soil using a 1mm sieve. The soil that passed through the sieve was designated as the small particle size treatment (Sm), and the soil that did not was designated as the large particle size treatment (La). As a control, we used field soil at room temperature (FS-control, also referred to as FS). We started 15 replicates of each treatment and sampled five replicates of each treatment at 2 weeks, 10 weeks, and 24 weeks post-inoculation. Because the mesocosms took ~ 1 month to dry out, there is no 2-week Dry treatment. We started an additional five replicates of the FS-control and Clay to assess initial population sizes and an additional 5 mesocosms of the FS-control for a long-term timepoint, which we sampled after 2.5 years. No water was added after the 24-week time point.

At each sampling time, we transferred the mesocosm contents to a sterile 50 mL Falcon tube and added 30 mL sterile H_2_0. We vortexed each tube on high for 1 minute and then placed the tubes on their sides and shook for 30 minutes (100 rpm) at 4°C. We then centrifuged the tubes at 400 × g for 8 min and transferred the supernatant to a new 50 mL tube, discarding the pellet of soil particles. To concentrate the bacteria, we then centrifuged at ~10,000 × g for 8 min and discarded the supernatant. We resuspended each pellet in 1 mL 0.85% NaCl. We took 10 μL of the supernatant to estimate population sizes using replicate dilution plating. We plated two replicates of each of the three dilutions (5, 6, 7) on TY plates, incubated them at 31°C for 36-48 hours, and counted the colonies. We averaged the two technical replicates as our estimate of population size. Lastly, we spun the remaining resuspended pellet at 10,000 × g for 5 minutes, removed the supernatant, and stored the bacterial pellets at −20°C until extraction. DNA was extracted using a Mo Bio Powersoil kit, following the standard instructions, except that we eluted the final DNA in 50 μL of RNase-free water.

We assessed rhizobia fitness in host nodules using the methods used by Burghardt et al. (2022) (36). Briefly, we grew six replicate pots with 10-12 plants of either *Medicago littoralis* R108 or *M. truncatula* A17, and immediately after planting, we inoculated each plant with 10^8^ cells of the same inoculum used to inoculate the soil mesocosms (Fig. 1). After 6 weeks, all the nodules were collected from each replicate pot, surface sterilized, and crushed with a pestle in sterile H_2_O. Debris and bacteroids were removed by centrifuging at 400 x g for 10 minutes and saving the supernatant. The undifferentiated rhizobia from the supernatant were then pelleted at high speed >16,000 x g for 5 minutes. Pellets were stored at −20°C until we extracted DNA using the UltraClean Microbial DNA Isolation Kit (no. 12224; Mo Bio Laboratories), following the standard instructions, except that we eluted the final DNA in 50 μL of RNase-free water.

### Sequencing to infer community composition and strain fitness

To estimate the strain frequencies in the nodule pools and in the soil samples, we followed the methods used by Burghardt et al. (2022) (36). The DNA extracted from each soil sample or nodule pool were sequenced on an Illumina HiSeq 2500 (NexteraXT libraries, 125 bp paired-end reads, and 3.6 to 9.4 million read pairs library^-1^) at the Joint Genome Institute. The reads were trimmed with TrimGalore! (v0.4.1; min length = 100bp, quality threshold = 30) and aligned to the *S. meliloti* USDA1106 genome (35) using bwa mem (v0.7.17; (37); NCBI BioProject: PRJNA388336). Median coverage per sample was 65× (range: 29× to 115×). We identified SNPs using FreeBayes(v1.2.0-2-g29c4002 (38), minimum read mapping quality of 30); and then estimated the frequency of each strain using HARP (39). We could not obtain strain frequency estimates from the two-week 4°C time point due to low DNA yield. We estimated strain relative fitness as the fold change in the frequency of each strain (q_x after selection_) in a nodule or soil community relative to the mean frequency of that strain across four sequencing replicates of the initial community (q_x initial inoculum_). Therefore, fitness is calculated as log_2_(q_x after selection_ /q_x initial inoculum_).

### Statistical analysis of population size, diversity, community composition, and potential host benefit

For the two univariate traits, population size and Shannon diversity, we ran ANOVA in R (‘lm’ and ‘anova’) using the R version 4.4.2 (2024-10-31). If we found significant interactions between the time and treatment variables, we ran separate analyses for each time point (or treatment). We used the Dunnett’s *post hoc* test using the multcomp package (40) to determine if each treatment (or timepoint) differed from field soil control (FS-control). If we did not find an interaction between variables, or when working with a single variable, we ran a Tukey HSD *post*-*hoc* test using the TukeyHSD() function. For population size, we used the log_10_-transformed Colony Forming Units (CFU) counts in each soil mesocosm to reduce the skew of the distribution (41). We calculated Shannon’s diversity, based on rhizobia frequency for each replicate using the ‘diversity’ function in the vegan package in R (42). For the multivariate trait, strain relative fitness, we performed Redundancy Analysis (RDA) using the vegan package, to assess how time, treatment, and their interaction (if relevant) affected strain fitness, and a permutation-based test (“anova” function, 1000 permutations) to assess the significance of each term. To assess whether specific treatments differed significantly from the FS-control, we reran the model to compare only the FS-control with the specific treatment.

We estimated the potential host benefit of the rhizobial community in the inoculum and each soil mesocosm at 24 weeks using data from a published single-strain inoculation experiment (6). We inferred the potential benefit of the soil community as in Burghardt et al. (2022) (36). Briefly, using a host-specific dataset, we took the summation of the per-strain frequency multiplied by the respective dry shoot weight and then divided that by the sum of the strain frequencies. Then, we scaled each host-specific dataset from zero to one, with one representing 100% occupancy by the most beneficial strain and zero representing 100% occupancy by the least beneficial strain. We used an ANOVA (‘lm’ and ‘anova’) with treatment as a predictor to analyze differences in potential host benefit across soil perturbations. We used Dunnett’s *post hoc* test to assess, first, whether the potential benefit of the community in the initial inoculum differed from that of the soil-selected rhizobial communities, and second, whether the potential benefit of the community in FS-control differed from that of the community after abiotic perturbations.

### Phenotypic fitness correlations and Genome-wide association

Pearson’s correlations were calculated using the cor.test() function in R. For the fitness correlations, we used the median strain relative fitness of each strain in their respective environment, either host or soil. For the correlations between plant benefit and rhizobial relative fitness in the soil, we used the aboveground biomass from a published single-strain inoculation experiment (data from Burghardt *et al*. 2018 (6)). We calculated all the correlations using the strain fitness values in the soil taken at 24 weeks.

The GWAS analysis was run with the previously filtered SNPs that had a MAF < 5% (6). We used a linear mixed model implemented in GEMMA (43) to determine the allelic effect sizes for rhizobia strain fitness in the soil or in the host. Additionally, to control for rhizobia population structure, we used the standardized kinship matrix created with GEMMA as a covariate. The variants were grouped in linkage disequilibrium groups (LD groups), based on non-random association of variants (34). We used likelihood ratio tests to calculate the P-values for each LD group. Because the rhizobia replicons have different population structures, the GWAS analysis was run separately for each replicon (34). The number of LD groups varied per replicon (2910 chromosomal, 12396 pSymA, and 17653 pSymB variants).

## RESULTS

### Temperature and salt most strongly affect population sizes, diversity and community composition

Rhizobial population sizes in most treatments showed similar dynamics as the FS-control; population size increased sharply during the first two weeks, then declined slightly (Fig. 2A). By contrast, at 4°C, the population growth was far slower, resulting in a lower population size than the FS-control (Time × Temperature P < 0.001, F= 59.958, Table S1A and Table S1B) and at 32°C, the population grew similar to the FS-control during the first 2 weeks, but then experienced the sharpest decline (Table S1B). Similar to population sizes, Shannon’s diversity of the rhizobial community was more strongly affected by temperature than by the other treatments, especially at the longest time points (Fig. 2B, Time × Treatment, F = 5.874, P < 0.001, Table S2A, Table S2B). Cold treatment at 24 weeks resulted in the lowest diversity, suggesting that even though growth is slow, selection is strong (Fig. 2B). Indeed, a handful of strains dominate in the 4°C environments (Fig. 2D).

**Figure 2:**
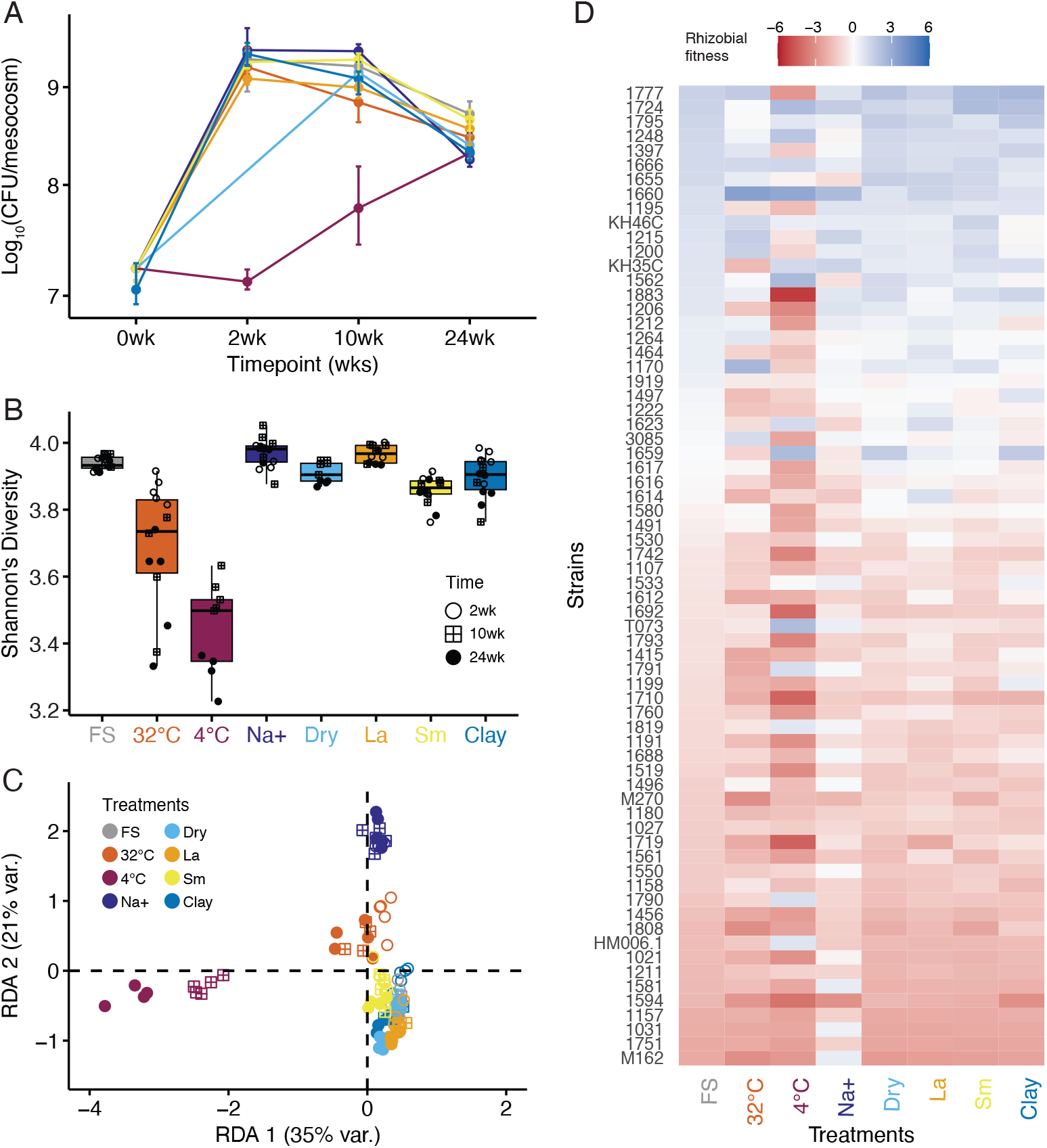
Abiotic treatments applied in the soil treatments affect population size, diversity, and rhizobial community composition. (A) Population growth dynamics across all soil treatments at three time points. We transformed the Colony Forming Units (CFU) values per mesocosm (tube) to a log scale (log_10_). We did not measure the population size of the Dry treatment at 2 weeks because the treatment had not yet been imposed. (B) Shannon diversity of the rhizobial community at three timepoints. (C) Redundancy Analysis (RDA) visualizing the variation in rhizobial relative fitness that soil treatments can explain over time (model = relative strain fitness ~ treatment*time). (D) Heatmap of median strain fitness values at 24 weeks. The soil treatments include field soil at room temperature (FS-control or FS), warm temperature (32°C), cold temperature (4°C), high salinity (Na+), drought (Dry), large particle size (La), small particle size (Sm), and clay. See Tables S1, S2, S3, and S4 for full statistical results.

The soil community composition (strain relative fitness) was strongly affected by the treatments (Treatment, P < 0.001, F = 89.76, Table S3B), with the magnitude of the effect shifting over time (Time × Treatment, P < 0.001, F = 5.33, Table S3B, Fig. 2C). Pairwise RDA models comparing each abiotic treatment to the FS-control indicate that the treatments 4°C, Na+, and 32°C explained the greatest proportion of the variation (Table S4) and thus drove more substantial shifts in rhizobial communities. The effects of 4°C on the community differed from those of 32°C, which showed similar effects to Na+ (Fig. 2C).

Most of our treatments were measured for a maximum of 24 weeks; however, since *S. meliloti* can survive in the soil for years (44), we sought to evaluate further the effect of long-term incubation on rhizobial diversity and community composition. To do this, we sampled some of the FS-control mesocosms 2.5 years after inoculation. We observed that time had a significant effect on both diversity (Table S5, Fig. S1A) and community composition (Table S6, Fig. S1B), with the largest shifts in both metrics occurring after 2.5 years. The 2.5-year sample had significantly lower diversity than earlier time points, which did not differ significantly from each other (Table S5B, Fig. S1A).

### Rhizobial fitness is correlated across soil conditions, but it is not correlated between soil and hosts

At 24 weeks, all treatment’s population sizes were similar, so we focus on this timepoint for our GWAS and fitness correlations (see Estimates in Table S1). We found positive correlations in strain fitness between each pair of soil environments (Fig. 3A, Fig. S2, Table S7). The pairwise correlations between most soil environments were relatively strong (R> 0.7, Table S7, Fig. 3C, Fig. S2), suggesting that rhizobia with high fitness in one soil environment condition also perform well in the other. The exceptions to these strong correlations were those involving 4°C (R ranges from 0.365 to 0.455, Fig. 3E), salinity (R ranges from 0.367 to 0.644), and 32°C (R ranges from 0.365 to 0.791). Overall, our results suggest rhizobia possess similar adaptations to succeed in multiple soil conditions.

**Figure 3:**
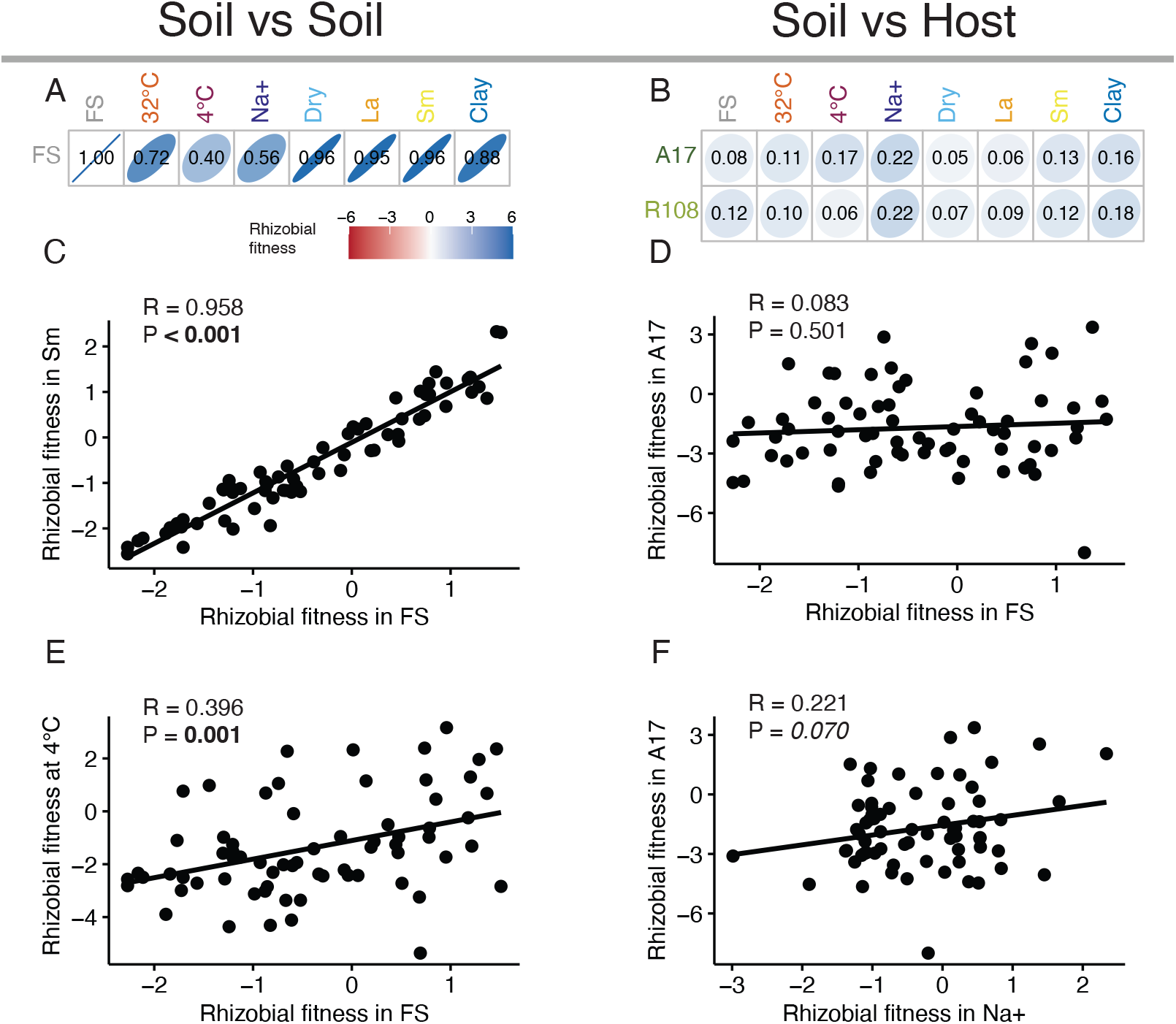
There is a positive correlation between the rhizobial strain fitness across soil treatments, but no correlation between the soil treatments and the host plants. Plot of Pearson correlation coefficients (R) between Field Soil control (FS) and the abiotic soil treatments (A) and between all the soil treatments and the two host plants, *Medicago truncatula* A17 and *Medicago littoralis* R108 (B). We found all significant positive correlations between soil environments, for instance, between FS and Small particle size (Sm) (C), and between FS and 4°C (E). In contrast, we found no significant correlations between field soil and host environments (e.g., FS and A17 hosts (D)), but salt soil additions led to marginally significant, weak correlations with A17 (F) and R108. The strain fitness measurements in the soil used in this analysis were calculated at 24 weeks. Each black dot represents a strain. See Table S7 and Table S8 for full results.

In contrast to the positive correlations across soil environments, we found that the rhizobial fitness in the soil was not correlated with the fitness inside hosts (R ranges from 0.046 to 0.221, all P > 0.05, Fig. 3B, 3D, 3F, Table S8). This suggests that the rhizobial adaptations in the bulk soil and in the plant are independent from each other.

To determine whether shared genetic variants are underlying the fitness correlations, we ran a GWAS on the rhizobial fitness in each soil and host environment at 24 weeks. We found that the strength of individual associations and was relatively weak—only six loci (5 on pSymA and 1 on pSymB) passed the Bonferroni-corrected threshold, revealing a complex genetic architecture where many loci of small effect influence strain fitness in the soil. The only soil treatments that had at least one locus statistically significantly associated with rhizobial fitness were 4°C and 32°C, and clay (Fig. S3, Table S9). Furthermore, none of the loci significantly affected rhizobial relative fitness in two environments, either when comparing across soil conditions (Fig. S4) or between hosts and soil treatments (Fig. S5).

### The environment during soil selection affects the abundance of high- and low-quality strains

Two complementary analyses revealed that soil selection often weakly increases the abundance of high-quality partners—with some exceptions. First, we examined correlations between strain relative fitness in the soil and plant shoot biomass in a single-strain inoculation, N-free conditions (Figure 4A, 4B). Most correlations were minimal and not significant (R ranged from −0.145 to 0.304, most P > 0.05, Table S10) except soil selection in 4°C and benefits to the A17 host, which were weakly positively correlated (R = 0.320, P = 0.013, Fig. 4A, Table S10). Second, we calculated the potential host benefit of each mesocosm’s strain community to ask if soil treatments enriched the rhizobial community for better quality strains (those providing higher shoot biomass). When using the initial inoculum as a comparison point, we observed that, for R108, most of the soil conditions, including the FS-control, were slightly enriched for high-quality partners, except for 4°C, which was enriched with low-quality partners. Similarly, for A17, most of the soil conditions are slightly enriched with high-quality partners except FS-control and Large particle size (La), that do not significantly enrich either high- or low-quality partners (Table S11A, Table S11B, Fig. S6). Furthermore, when using the FS-control as a baseline comparison against abiotic perturbations, we observed that, for A17, both 32°C and 4°C conditions enriched for better partners (Fig. 4C, P < 0.001, Table S12A, Table S12B). In contrast, for R108, 4°C selected for lower-quality strains, while selection under 32°C, small particle size soil (Sm), and clay enriched the soil for better R108 partners (Fig. 4D, P < 0.01, Table S12A, Table S12B). These results suggest that soil selection can alter the rhizobial populations in ways that significantly influence the relative abundance of high- and low-quality strains. However, the effects are host-dependent and relatively modest.

**Figure 4:**
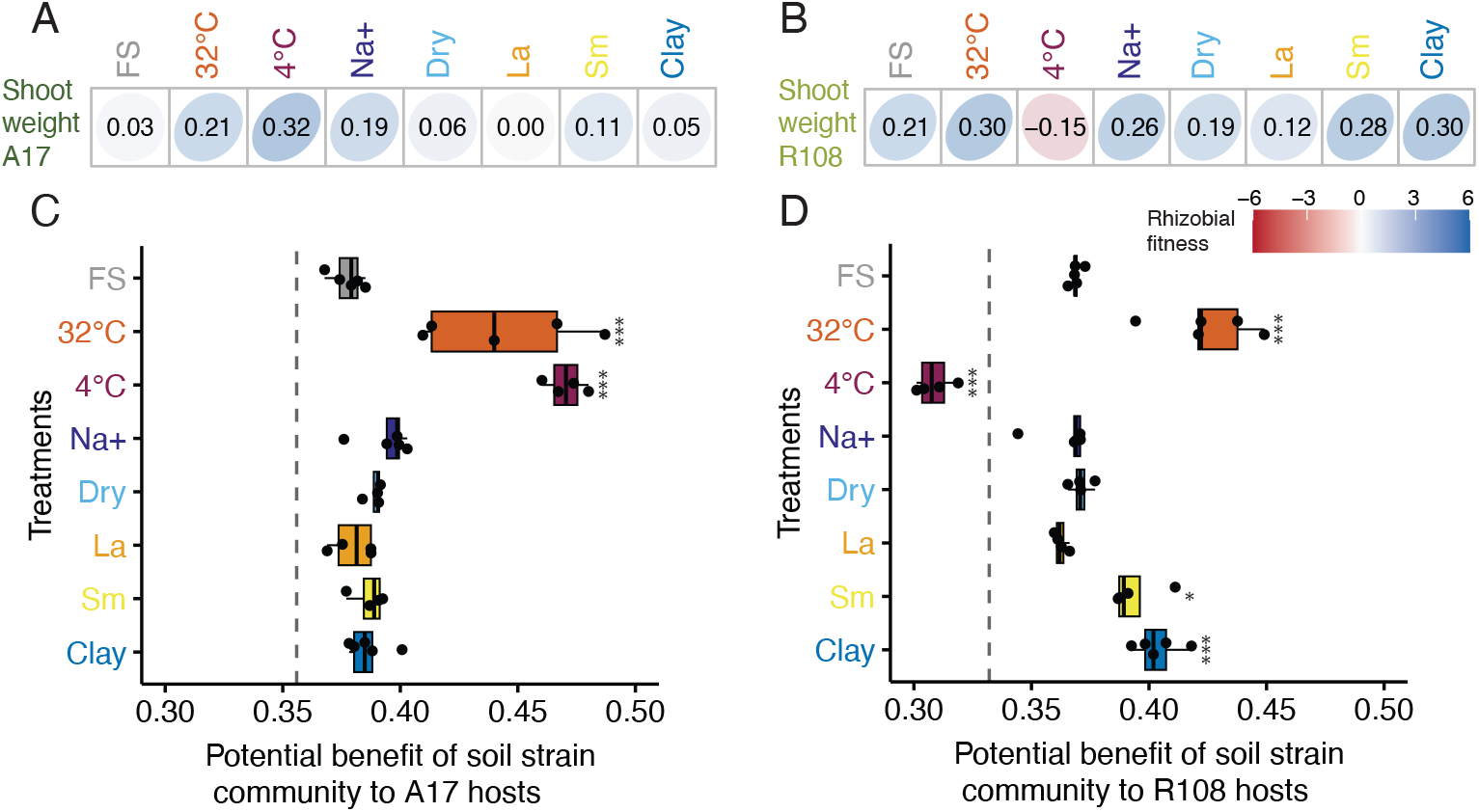
Temperature changes can alter the quantity of high-quality and low-quality partners, depending on the host. Plots of the Pearson correlation coefficients between rhizobial relative fitness in the soil at 24 weeks and the shoot biomass of *Medicago truncatula* A17 (A) and correlations between rhizobial relative fitness in the soil at 24 weeks and the shoot biomass of *Medicago littoralis* R108 (B). The potential benefit of the soil strain community to A17 hosts (C) and R108 hosts (D). The potential benefit was calculated as a summation of the per-strain frequency multiplied by the dry shoot weight, and each host-specific dataset was scaled from zero (soil community is entirely composed of the least beneficial strain) to one (soil community is entirely made up of the most helpful strain). The dashed line is the estimated benefit provided by the initial community. We used one-way ANOVA followed by Dunnett *post hoc* test. All treatments were compared to the FS-control (FS), and significantly different samples are denoted as follows: ***, P < 0.001; *, P < 0.05. See Table S10 and Table S12 for full statistical results. See Table S11 and Figure S6 for comparisons of each treatment to the initial community.

## DISCUSSION

Understanding the selection rhizobia undergo when living in the soil and after they have established themselves in the plant’s nodules will provide a more comprehensive understanding of rhizobial evolution in agricultural ecosystems. Here, we leveraged the natural genetic diversity of 68 *S. meliloti* strains to assess the effect of soil selection under multiple abiotic perturbations and in two *Medicago* hosts. We observed that 1) temperature perturbations had significant effects on population size and rhizobial diversity; additionally, temperature and salinity treatments had strong effects on rhizobial community composition; 2) There was a positive correlation between the rhizobial relative fitness across different soil treatments, but there were no correlations between the host and soil treatments. Furthermore, we found few strong associations between genetic variants and rhizobial fitness, consistent with soil fitness being a complex genetic trait. 3) Intriguingly, temperature treatments seem to have an indirect effect on the selection of rhizobia that provide higher or lower plant benefit with the direction of the selection depending on the host genotype. Overall, our results indicate that rhizobia exhibit common adaptations across various soil conditions, yet follow an independent evolutionary trajectory as compared to when living in association with their host plant.

We observed that temperature and saline treatments had the greatest effects on the community metrics we assessed, particularly cold perturbations caused the most dramatic decrease in diversity, and the strongest shift in community composition, which could be explained by only a few strains of rhizobia having a disproportionate ability to grow at 4°C (45–48). We also observed that most of the genetic variants that were significantly associated with rhizobium relative fitness across all the soil treatments in our GWAS study at 24 weeks (four of the six genetic variants that crossed the Bonferroni threshold) were found in the soil at 4°C and are on PsymA. These results were unexpected for two reasons. First, pSymA is primarily known as the symbiotic plasmid and is not typically considered to be involved in adaptation to abiotic conditions, although it does possess genes that could confer differential fitness in the soil (49–51). Second, none of the genes we identified have been previously reported as important for microbial fitness in cold conditions. For instance, RepA is involved in regulating plasmid partitioning and stability in multiple soil bacteria (52), genes containing PAS-domains can be associated with quorum sensing and motility (53), and genes encoding mechanosensitive ion channels are associated with protection against osmotic stressors (54). Because the correlation between field soil fitness and fitness in the cold is weakly positive, these genes could also be important for fitness in the soil more generally.

Our results showed that, in most of our soil treatments, strains with higher fitness in one soil environment tend to have higher fitness in other soil environments, suggesting the presence of cross-protective strategies, where adaptation to one stressor provides adaptation to another, as the result of evolution to predictable or co-occurring perturbations in their habitat (55–57). An exception to these strong positive correlations between rhizobial relative fitness across environments was the weak correlations observed between 4°C and other soil perturbations, suggesting that cold selection has strong, environmentally specific effects (58, 59). These weak correlations, particularly between 4°C and 32°C could have implications for rhizobial populations in nature because diurnal and seasonal temperature fluctuations are common in agricultural and natural ecosystems. While we used static conditions, our results suggest that seasonal shifts between hot and cold temperatures can differentially increase the strain growth rates across the year, which we speculate could have a cascading effect during periods of root growth and nodule formation (60). While we found an overall signal of soil fitness alignment across all abiotic perturbations, further studies in natural ecosystems are needed to assess the implications of these strong and weak fitness correlations on rhizobial populations under fluctuating environments.

Rhizobial fitness in the soil was not correlated with rhizobial fitness in plants. We expected that rhizobia would co-opt the protective mechanisms they use to face abiotic stresses in the soil, such as osmotic and oxidative stressors (61–64), to cope with the perturbations they encounter during root infection and nodule formation (31, 52). Our results, however, suggest that the nodule and the soil are substantively different environments (65). Our results align with studies using mutant libraries that have compared rhizobial competitive fitness in the rhizosphere, an out-of-host habitat, versus the nodule, since some mutants outcompeted in the rhizosphere did not necessarily have impaired competitive fitness in the nodule (26, 66). The minimal correlations between adaptive traits in both habitats could indicate that selection favors the evolution of independent genetic pathways underlying those traits, thereby avoiding negative trade-offs that could constrain adaptation in either trait. Our study provides novel insights into the selection that a symbiont faces during its outside-the-host stage; however, further studies are needed to assess whether symbionts in other facultative mutualisms exhibit similarly weak correlations when they are inside and outside the hosts.

Our results suggest that, in many cases, the rhizobial reproductive success in the soil is independent of whether the strain benefits the plant. These results have implications for the maintenance of variation in partner quality (67). A growing body of literature suggests that in a multi-strain context, the plant rewards and thereby selects more beneficial strains (6, 32, 68), therefore reducing genetic diversity (22, 69, 70). To understand what counteracts positive selection (7), studies have focused on plants as drivers of the maintenance of genetic diversity (32, 71, 72). We propose the bulk soil as another environment to maintain variation. Our observations suggest that genes that affect plant benefit may evolve through drift when rhizobia are in the soil, rather than through trade-offs, as proposed by Heath & Stinchcombe, (2014) (7). Moreover, we showed two unexpected results. High temperatures consistently increased the relative fitness beneficial rhizobia, while cold temperatures either increased or decreased the relative fitness of beneficial bacteria, depending on the host genotype. While there may be environmental stressors inside the host that are similar to our soil perturbations, an alternative possibility is that these results are an example of coincidental evolution, where abiotic factors outside the host indirectly select for a host-associated trait i.e. high temperatures select for *Vibrio fischeri* with better colonization and an increased bioluminescence in their squid host (73), or for *Serratia* that are more pathogenic for wood tiger moth (74). Our findings indicate that the selection outside the host has implications for the functioning of the mutualism, and therefore, we highlight the importance of evaluating the evolutionary dynamics of a symbiont during its various life-history stages (75, 76).

Our work demonstrates that, under static treatments, the soil and host do affect the selection of rhizobial strains. However, to understand how these factors interact, it is necessary to assess natural genetic variation in the field (77). Also, intriguingly, temperature treatments in the soil appear to be selecting for either high- or low-quality rhizobia; however, further validation is necessary to determine if this has implications for rhizobial colonization of plants, such as by using hosts that experience multiple seasons during their lifetime, like perennial legumes. Finally, since we are working with soil as a substrate, and testing multiple abiotic perturbations, our study has found candidate genes, particularly in pSymA, that have not been described before as important for rhizobial fitness, highlighting the importance of replicating natural conditions, because rhizobia exists in conditions far from the “optimal” in-vitro used in most assessments (50, 77). Understanding how selection occurs in a diverse community of rhizobia across their life history provides insight into what happens in a real agricultural ecosystem, where multiple factors, such as host genotype and soil abiotic stressors, interact (27).

## Acknowledgment

This work was supported by the National Science Foundation grants IOS-1724993,1856744, 2243817, and 2243819 to Burghardt and Tiffin and completed through computing resources provided by the Minnesota Supercomputing Institute (MSI) at the University of Minnesota and sequencing support from the Department of Energy Community Sequencing Project #503446. LTB’s work was supported by the USDA National Institute of Food and Agriculture and Hatch Appropriations under Project #PEN04760 and Accession #1025611. We thank additional members of the Tiffin Lab for their help in plant processing.

## Data availability

All data and analysis will be made publicly available in a repository upon publication. Current analysis can be found at: https://github.com/mariale4915/Soil_Mesocosm_Manuscript

