## Supplementary Material for "Rhizobia independently adapt to soil and legume host environments, but soil conditions influence the abundance of high-quality partners"

**Table S1:** The abiotic treatments (Trt) affect the population size across timepoints (Time). The population size is represented as $log_{10}$(CFU/mesocosm). (A) Two-way ANOVA Full Model. (B) Dunnett's *post hoc* test compared population size in field soil control (FS-control, also referred to as FS) with each soil perturbation (listed under Comparison) within each timepoint. Df (Degrees of freedom), model sum of squares (Sum sq), P (P-value), F value (F-statistics), Estimate (Estimated differences in means).

(A)

| Term | Df | Sum Sq | F value | P |
|---|---|---|---|---|
| Time | 3 | 70.310 | 1886.231 | **< 0.001** |
| Trt | 7 | 15.672 | 180.183 | **< 0.001** |
| Time:Trt | 20 | 14.900 | 59.958 | **< 0.001** |
| Residuals | 113 | 1.404 | NA | |

(B)

| Comparison | 2 wk | | 10 wk | | 24 wk | |
|---|---|---|---|---|---|---|
| | **Estimate** | **P** | **Estimate** | **P** | **Estimate** | **P** |
| 32°C - FS | 0.054 | 0.972 | -0.446 | **< 0.001** | -0.262 | **0.004** |
| 4°C - FS | -2.093 | **< 0.001** | -1.459 | **< 0.001** | -0.385 | **< 0.001** |
| Na+ - FS | 0.223 | *0.068* | 0.143 | 0.367 | -0.458 | **< 0.001** |
| La - FS | -0.124 | 0.518 | -0.247 | **0.028** | -0.179 | **0.010** |
| Sm - FS | 0.118 | 0.566 | 0.042 | 0.995 | -0.090 | 0.388 |
| Clay - FS | 0.136 | 0.429 | -0.161 | 0.257 | -0.419 | **< 0.001** |
| Dry - FS | | | -0.071 | 0.917 | -0.347 | **< 0.001** |

**Table S2**: Temperature treatment has a significant effect in the rhizobial diversity at later timepoints. (A) Two-way ANOVA, full model. (B) Dunnett's *post hoc* test on Shannon diversity comparing the field soil control (FS) to all the others abiotic treatments (Comparison) within each timepoint. We were unable to calculate the frequency of the rhizobial strains at 4 °C in the 2-week time point due to a low DNA yield. We also did not calculate Dry at 2 weeks because the tubes took around 4 weeks to dry out.

(A)

| **Term** | **Df** | **Sum Sq** | **F value** | **P** |
|---|---|---|---|---|
| Trt | 7 | 2.429 | 84.630 | **< 0.001** |
| Time | 2 | 0.107 | 13.065 | **< 0.001** |
| Trt:Time | 12 | 0.289 | 5.874 | **< 0.001** |
| Residuals | 80 | 0.328 | NA | |

(B)

| **Comparison** | **2wk** | | **10wk** | | **24wk** | |
|---|---|---|---|---|---|---|
| | **Estimate** | **P** | **Estimate** | **P** | **Estimate** | **P** |
| 32°C - FS | -0.067 | *0.052* | -0.332 | **< 0.001** | -0.376 | **< 0.001** |
| Clay - FS | 0.019 | 0.914 | -0.073 | 0.517 | -0.074 | 0.433 |
| La - FS | 0.046 | 0.393 | 0.027 | 0.989 | 0.008 | 1.000 |
| Na+ - FS | 0.026 | 0.745 | 0.028 | 0.985 | 0.036 | 0.942 |
| Sm - FS | -0.069 | **0.043** | -0.088 | 0.274 | -0.093 | 0.259 |
| 4°C - FS | | | -0.404 | **< 0.001** | -0.625 | **< 0.001** |
| Dry - FS | | | -0.017 | 0.999 | -0.059 | 0.712 |

**Table S3:** Redundancy analysis reveals a significant interaction of time and treatment (Trt) in the rhizobial community composition in the soil. (A) Variance partitioning of rhizobial strain fitness explained by Time and Trt. The component "Constrained" represents the variance explained by the explanatory variables Time and Treatment, whereas the "Unconstrained" component is the variance not explained by those explanatory variables. The column "Inertia" represents the total variance of the data explained by the constrained or unconstrained component while the "Proportion" column represents the proportion of the total variation explained by both components. (B) Permutation test results for the Full RDA model.

(A)

| **Component** | **Inertia** | **Proportion** |
|---|---|---|
| Total | 68 | 1 |
| Constrained | 61.257 | 0.901 |
| Unconstrained | 6.743 | 0.11 |

(B)

| **Term** | **Df** | **Prop.Var** | **F value** | **P** |
|---|---|---|---|---|
| **$R^2$adj =** | 0.875 | | | |
| Trt | 7 | 0.779 | 89.76 | **0.001** |
| Time | 2 | 0.043 | 17.2 | **0.001** |
| Trt:Time | 12 | 0.079 | 5.33 | **0.001** |
| Residual | 80 | 0.099 | | |

**Table S4**: Cold, salinity, and warm treatments explained the strongest community shifts. Pairwise comparisons were performed using permutation test on RDA models that included Treatment and Time. Each comparison represents the contrast between the field soil control (FS) vs other soil treatments.

| Comparison | $R^2$.adj | Term | Df | F value | P |
|---|---|---|---|---|---|
| 4°C - FS | 0.872 | **Trt** | 1 | 120.06 | **0.001** |
| | | **Time** | 2 | 12.91 | **0.001** |
| | | **Trt:Time** | 1 | 14.37 | **0.001** |
| | | Residual | 19 | | |
| Na+ - FS | 0.704 | **Trt** | 1 | 57.3 | **0.001** |
| | | **Time** | 2 | 5.36 | **0.001** |
| | | **Trt:Time** | 2 | 3.06 | **0.02** |
| | | Residual | 24 | | |
| 32°C - FS | 0.609 | **Trt** | 1 | 30.2 | **0.001** |
| | | **Time** | 2 | 4.65 | **0.002** |
| | | **Trt:Time** | 2 | 4.56 | **0.002** |
| | | Residual | 23 | | |
| Clay - FS | 0.584 | **Trt** | 1 | 20.31 | **0.001** |
| | | **Time** | 2 | 8.35 | **0.001** |
| | | **Trt:Time** | 2 | 3.64 | **0.002** |
| | | Residual | 23 | | |
| La - FS | 0.469 | **Trt** | 1 | 11.72 | **0.001** |
| | | **Time** | 2 | 6.79 | **0.001** |
| | | Trt:Time | 2 | 1.34 | 0.198 |
| | | Residual | 21 | | |
| Dry - FS | 0.443 | **Trt** | 1 | 6.47 | **0.001** |
| | | **Time** | 2 | 6.43 | **0.001** |
| | | **Trt:Time** | 1 | 2.94 | **0.006** |
| | | Residual | 19 | | |
| Sm - FS | 0.443 | **Trt** | 1 | 11.39 | **0.001** |
| | | **Time** | 2 | 6.9 | **0.001** |
| | | Trt:Time | 2 | 1.05 | 0.396 |
| | | Residual | 23 | | |

**Table S5:** Shannon diversity in the field soil control (FS) is influenced by the time. (A) Full ANOVA model. (B) Tukey HSD *post-hoc* test used to compare diversity across timepoints (Comparison). Diff (Difference between the means of two groups)

(A)

| **Term** | **Df** | **Sum Sq** | **F value** | **P** |
|---|---|---|---|---|
| Time | 3 | 0.071 | 41.545 | **< 0.001** |
| Residuals | 16 | 0.009 | NA | |

(B)

| **Comparison** | **Diff** | **P** |
|---|---|---|
| 2wk-10wk | -0.025 | 0.376 |
| 2wk-24wk | -0.013 | 0.839 |
| 2wk-2.5y | 0.124 | **< 0.001** |
| 10wk-24wk | -0.012 | 0.841 |
| 10wk-2.5y | -0.149 | **< 0.001** |
| 24wk-2.5y | 0.136 | **< 0.001** |

**Table S6:** Redundancy analysis reveals a significant effect of time on the rhizobial community composition in the field soil (FS-control). (A) Variance partitioning of rhizobial strain fitness explained by Time. Shown are the total variance (Inertia) and the proportion of variance explained by the constrained component (Time), and the unconstrained (residual) component. (B) Permutation test results for the RDA model.

(A)

| **Component** | **Inertia** | **Proportion** |
|---|---|---|
| Total | 68 | 1 |
| Constrained | 41.699 | 0.613 |
| Unconstrained | 26.301 | 0.387 |

(B)

| **Term** | **Df** | **Prop.Var** | **F value** | **P** |
|---|---|---|---|---|
| **R2.adj =** | 0.541 | | | |
| Time | 3 | 0.613 | 8.456 | **0.001** |
| Residual | 16 | 0.387 | | |

**Table S7:** Pearson correlation coefficients (R) show that the rhizobial relative fitness at 24 weeks is positively correlated across soil treatments.

| Comparison | R | P |
|---|---|---|
| FS vs 4°C | 0.396 | **0.001** |
| FS vs 32°C | 0.723 | **<0.001** |
| FS vs Na+ | 0.563 | **<0.001** |
| FS vs Clay | 0.883 | **<0.001** |
| FS vs Sm | 0.958 | **<0.001** |
| FS vs La | 0.949 | **<0.001** |
| FS vs Dry | 0.959 | **<0.001** |
| 4°C vs 32°C | 0.365 | **0.002** |
| 4°C vs Na+ | 0.367 | **0.002** |
| 4°C vs Clay | 0.387 | **0.001** |
| 4°C vs Sm | 0.444 | **<0.001** |
| 4°C vs La | 0.455 | **<0.001** |
| 4°C vs Dry | 0.437 | **<0.001** |
| 32°C vs Na+ | 0.591 | **<0.001** |
| 32°C vs Clay | 0.558 | **<0.001** |
| 32°C vs Sm | 0.791 | **<0.001** |
| 32°C vs La | 0.671 | **<0.001** |
| 32°C vs Dry | 0.707 | **<0.001** |
| Na+ vs Clay | 0.553 | **<0.001** |
| Na+ vs Sm | 0.644 | **<0.001** |
| Na+ vs La | 0.442 | **<0.001** |
| Clay vs Dry | 0.897 | **<0.001** |
| Clay vs Sm | 0.865 | **<0.001** |
| Clay vs La | 0.841 | **<0.001** |
| Clay vs Dry | 0.897 | **<0.001** |
| Sm vs La | 0.895 | **<0.001** |
| Sm vs Dry | 0.950 | **<0.001** |
| La vs Dry | 0.932 | **<0.001** |

**Table S8:** Pearson correlation coefficients (R) show no significant correlations between rhizobial strain relative fitness in the soil at 24 weeks and rhizobial strain relative fitness in nodules of the hosts *Medicago truncatula* A17 or *M. littoralis* R108.

| Comparison | R | P |
|---|---|---|
| A17 - FS | 0.083 | 0.501 |
| A17 - 4°C | 0.166 | 0.175 |
| A17 - 32°C | 0.112 | 0.364 |
| A17 - Na+ | 0.221 | *0.070* |
| A17 - Clay | 0.162 | 0.188 |
| A17 - Sm | 0.125 | 0.308 |
| A17 - La | 0.064 | 0.602 |
| A17 - Dry | 0.046 | 0.707 |
| R108 - FS | 0.121 | 0.326 |
| R108 - 4°C | 0.059 | 0.634 |
| R108 - 32°C | 0.103 | 0.404 |
| R108 - Na+ | 0.219 | *0.073* |
| R108 - Clay | 0.181 | 0.140 |
| R108 - Sm | 0.118 | 0.337 |
| R108 - La | 0.086 | 0.484 |
| R108 - Dry | 0.072 | 0.559 |

**Table S9:** The LD groups that crossed the Bonferroni threshold in the Genome Wide Association Study were limited to three soil abiotic perturbations. The Bonferroni threshold differed per Replicon because the GWAS was run independently in each replicon: Chromosome ($1.718*10^{-05}$), pSymA ($4.036*10^{-06}$), and pSymB ($2.832*10^{-06}$). Replicon (Replicon where the LD group is located), Beta (allelic effect size), P (P-value, significance), Group size (LD group size that shows the number of genes in each group), locus tag (the NCBI locus tag), Product (putative product).

| Trt | Replicon | Beta | P | Group size | Locus_tag | Product |
|---|---|---|---|---|---|---|
| 32°C | PsymB | 0.915 | $2.75*10^{-06}$ | 1 | CDO30_RS28455 | FAD-dependent oxidoreductase; protein_id=WP_015456630.1 |
| 4°C | PsymA | 1.34 | $1.39*10^{-06}$ | 1 | CDO30_RS21590 | PAS domain S-box protein; protein_id=WP_013845133.1 |
| Clay | PsymA | 0.760 | $1.4*10^{-06}$ | 1 | CDO30_RS19900 | hypothetical protein; protein_id=WP_010967343.1 |
| 4°C | PsymA | 1.447 | $2.35*10^{-06}$ | 1 | CDO30_RS21920 | repA, plasmid partitioning protein RepA; protein_id=WP_010968237.1 |
| 4°C | PsymA | 1.217 | $1.61*10^{-06}$ | 1 | CDO30_RS21920 | repA, plasmid partitioning protein RepA; protein_id=WP_010968237.1 |
| 4°C | PsymA | 1.573 | $1.74*10^{-06}$ | 1 | CDO30_RS21395 | mechanosensitive ion channel family protein; protein_id=WP_088239352.1 |

**Table S10:** Pearson correlation coefficients (R) show that there are only weak correlations between rhizobial fitness in the soil at 24 weeks and the rhizobial plant benefit to A17 (weight_A17) and R108 (weight_R108).

| **Comparison** | **R** | **P** |
|---|---|---|
| weight_A17 vs FS | 0.029 | 0.828 |
| weight_A17 vs 4°C | 0.320 | **0.013** |
| weight_A17 vs 32°C | 0.210 | 0.111 |
| weight_A17 vs Na+ | 0.191 | 0.147 |
| weight_A17 vs Clay | 0.053 | 0.688 |
| weight_A17 vs Sm | 0.107 | 0.419 |
| weight_A17 vs La | -0.001 | 0.996 |
| weight_A17 vs Dry | 0.056 | 0.676 |
| weight_R108 vs FS | 0.207 | 0.211 |
| weight_R108 vs 4°C | -0.145 | 0.385 |
| weight_R108 vs 32°C | 0.302 | *0.066* |
| weight_R108 vs Na+ | 0.258 | 0.118 |
| weight_R108 vs Clay | 0.304 | *0.063* |
| weight_R108 vs Sm | 0.279 | *0.090* |
| weight_R108 vs La | 0.121 | 0.470 |
| weight_R108 vs Dry | 0.186 | 0.262 |

**Table S11:** The potential benefits of the rhizobial community for hosts differ from the initial community after selection in the soil for 24 weeks in many cases, and the strength and direction depend on the host plant and soil treatments. (A) Results from separate ANOVA models for each host genotype with treatment as the predictor and potential benefit as the outcome. (B) Results of Dunnett's *post hoc* test run on each model in (A) that specifically compares the potential benefit of the initial inoculum (Initial) vs each soil treatment.

(A)

| **Host** | **Term** | **Df** | **Sum Sq** | **F value** | **P** |
|---|---|---|---|---|---|
| A17 | Trt | 8 | 0.043 | 27.359 | **< 0.001** |
| A17 | Residuals | 31 | 0.006 | | |
| R108 | Trt | 8 | 0.044 | 51.355 | **< 0.001** |
| R108 | Residuals | 31 | 0.003 | | |

(B)

| **Host** | **Comparison** | **Estimate** | **P** |
|---|---|---|---|
| A17 | 32°C - Initial | 0.087 | **< 0.001** |
| A17 | 4°C - Initial | 0.114 | **< 0.001** |
| A17 | Clay - Initial | 0.031 | **0.018** |
| A17 | Dry - Initial | 0.033 | **0.014** |
| A17 | FS - Initial | 0.022 | 0.143 |
| A17 | La - Initial | 0.024 | 0.116 |
| A17 | Na+ - Initial | 0.038 | **0.002** |
| A17 | Sm - Initial | 0.031 | **0.024** |
| R108 | 32°C - Initial | 0.093 | **< 0.001** |
| R108 | 4°C - Initial | -0.023 | **0.020** |
| R108 | Clay - Initial | 0.072 | **< 0.001** |
| R108 | Dry - Initial | 0.039 | **< 0.001** |
| R108 | FS - Initial | 0.037 | **< 0.001** |
| R108 | La - Initial | 0.030 | **0.002** |
| R108 | Na+ - Initial | 0.032 | **< 0.001** |
| R108 | Sm - Initial | 0.062 | **< 0.001** |

**Table S12**: The potential benefits of the rhizobial community for hosts differ from the community of the field soil control (FS-control) after selection in the soil for 24 weeks, and the strength and direction depend on the host plant and soil abiotic perturbations. (A) Results from separate ANOVA models for each host genotype with treatment as the predictor and potential benefit as the outcome. (B) Results of Dunnett's *post hoc* test run on each model in (A) that specifically compares the potential benefit of the FS-control (FS) vs each soil treatment.

(A)

| Host | Term | Df | Sum Sq | F value | P |
|---|---|---|---|---|---|
| A17 | Trt | 7 | 0.035 | 23.013 | **< 0.001** |
| | Residuals | 28 | 0.006 | | |
| R108 | Trt | 7 | 0.037 | 44.717 | **< 0.001** |
| | Residuals | 28 | 0.003 | | |

(B)

| Host | Comparison | Estimate | P |
|---|---|---|---|
| A17 | 32°C - FS | 0.066 | **< 0.001** |
| A17 | 4°C - FS | 0.093 | **< 0.001** |
| A17 | Clay - FS | 0.009 | 0.891 |
| A17 | Dry - FS | 0.012 | 0.768 |
| A17 | La - FS | 0.002 | 1.000 |
| A17 | Na+ - FS | 0.017 | 0.358 |
| A17 | Sm - FS | 0.009 | 0.899 |
| R108 | 32°C - FS | 0.056 | **< 0.001** |
| R108 | 4°C - FS | -0.060 | **< 0.001** |
| R108 | Clay - FS | 0.035 | **< 0.001** |
| R108 | Dry - FS | 0.002 | 1.000 |
| R108 | La - FS | -0.006 | 0.918 |
| R108 | Na+ - FS | -0.004 | 0.984 |
| R108 | Sm - FS | 0.025 | **0.010** |

**Figure S1:** Time alters rhizobial diversity and strain community composition in the field soil control (FS). (A) Shannon diversity decreases after 2.5 years, while there are no significant differences in diversity between the three initial timepoints (See Table S5). (B) Time contributed to variation in the rhizobial community composition in soil (See Table S6).

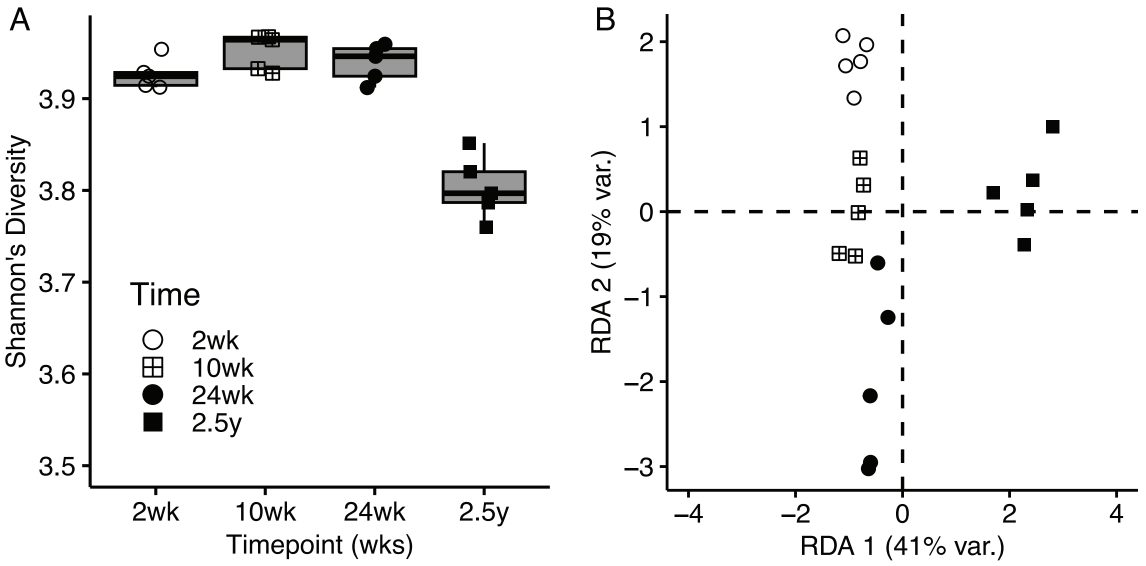

**Figure S2**: Strain relative fitness correlation coefficients (R) across all pairwise soil treatments at the 24-week timepoint are positive and significant. See Table S7 for R and P calculated for each Pearson correlation, and the color scale indicates the strength and direction of R.

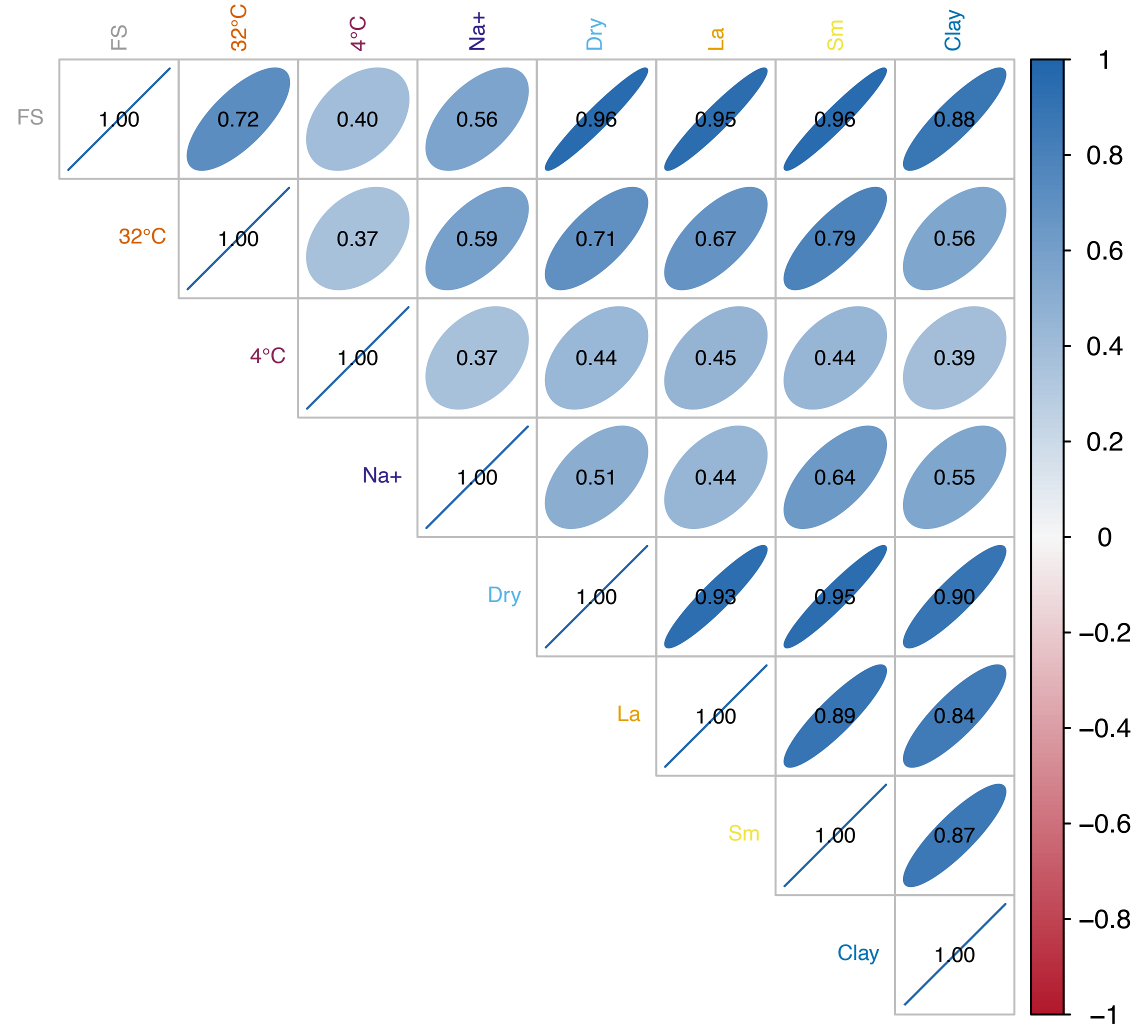

**Figure S3:** Rhizobial fitness in the soil is a complex trait with few LD groups showing statistical significance. Manhattan plot summarizing results of Genome-Wide Association (GWA) studies on rhizobial fitness in the seven soil environments tested. The x-axis shows the location of the LD group across the three replicons of the *Sinorhizobium meliloti* genome, and the y-axis shows the -log$_{10}$(P) of the strength of the association between strain fitness and genetic variation in each group. Colored dots indicate LD groups whose P-value (P) crossed the replicon-specific Bonferroni threshold (grey dashed lines). Significant associations were found in 4°C (purple circles), in 32°C (orange circles), or in clay (blue circles). See Table S9 for more information. The size of the circle was calculated by multiplying the absolute value of the allelic effect size by $10^5$

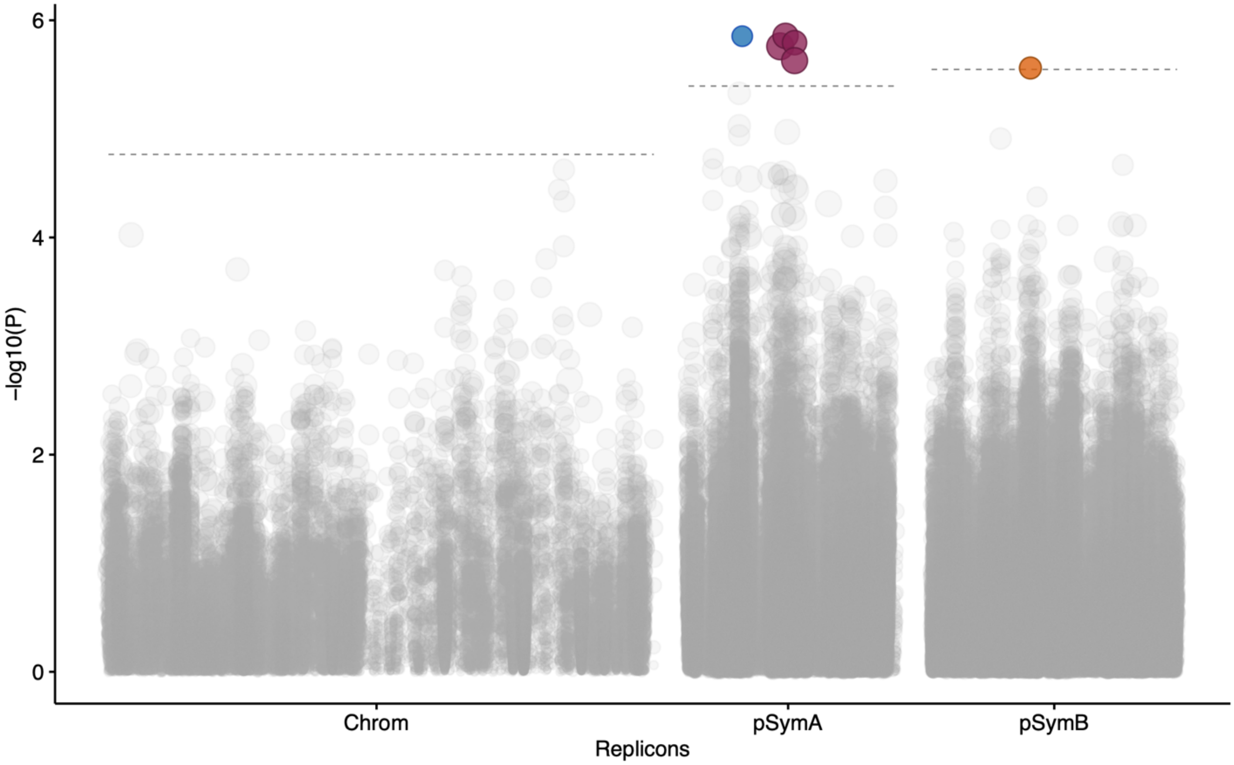

**Figure S4:** Pairwise comparisons between the allelic effect sizes (or beta values) of individual LD groups across all the soil conditions evaluated. Only 4°C (purple circle), 32°C (orange circle), and clay (blue circle) had LD groups that showed statistically significant association with rhizobia relative fitness (P-value crossed the Bonferroni threshold) in individual treatments. None of the LD groups were significant for more than one condition. The grey and transparent circles were not significant for any of the treatments.

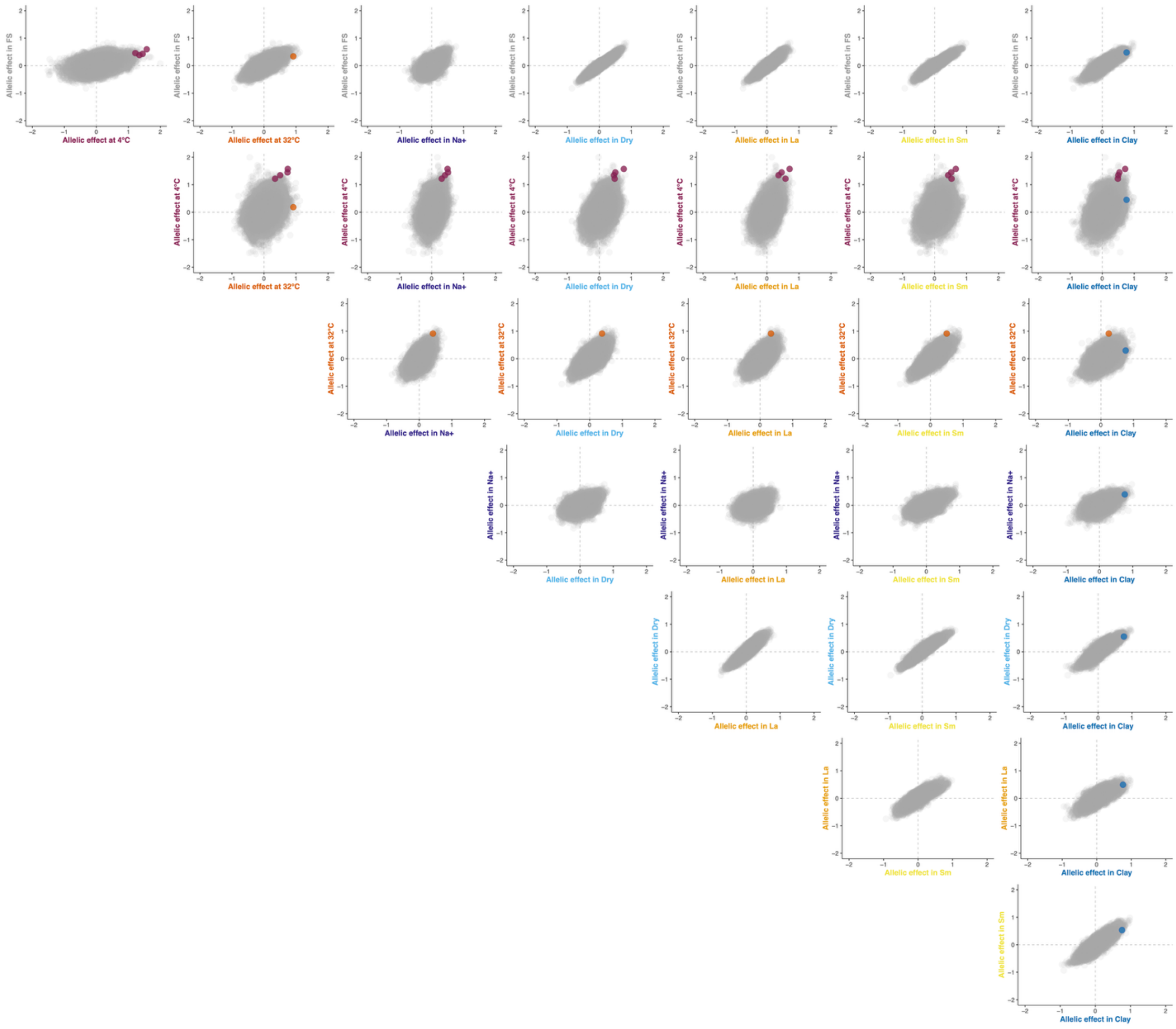

**Figure S5**: Pair-wise comparisons between the allelic effect sizes (or beta values) of individual LD groups between all the soil conditions and the two hosts, *Medicago truncatula* A17 and *Medicago* R108. The LD groups that show statistically significant association with rhizobia relative fitness (P-values cross the Bonferroni threshold) in either the host or the soil conditions are represented by circles that are either dark green (A17), light green (R108), purple (4°C), orange (32°C), and blue (Clay). None of the LD groups were significant for more than one condition. The grey and transparent circles were not significant for any of the treatments.

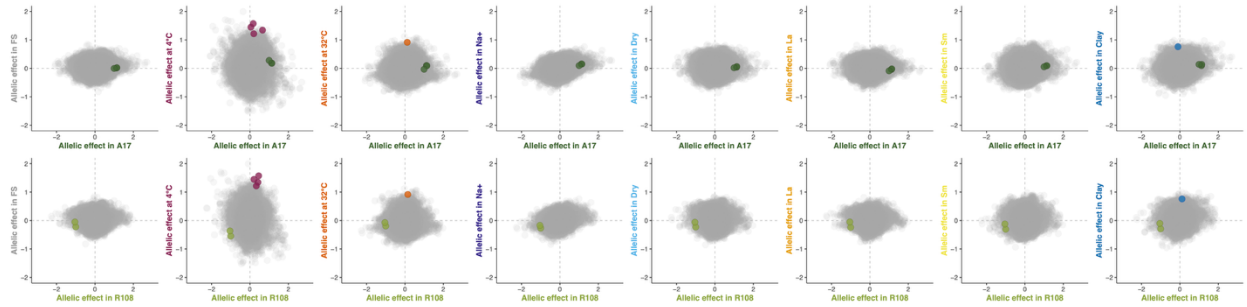

**Figures S6:** The rhizobial community selected by most soil treatments at 24 weeks has a higher or lower potential benefit than the initial inoculum, depending on the host plant, (A) *Medicago truncatula* A17, and (B) *Medicago littoralis* R108. We run a Dunnett's *post hoc* test comparing the potential benefit of the rhizobial community in the inoculum (Initial) vs the rhizobial community after each soil treatment. Samples are considered statistically significant when compared with the initial inoculum if: ***, P < 0.001; **, P < 0.01; *, P < 0.05. See Table S11B to see the Estimate, and P-value associated with this figure.

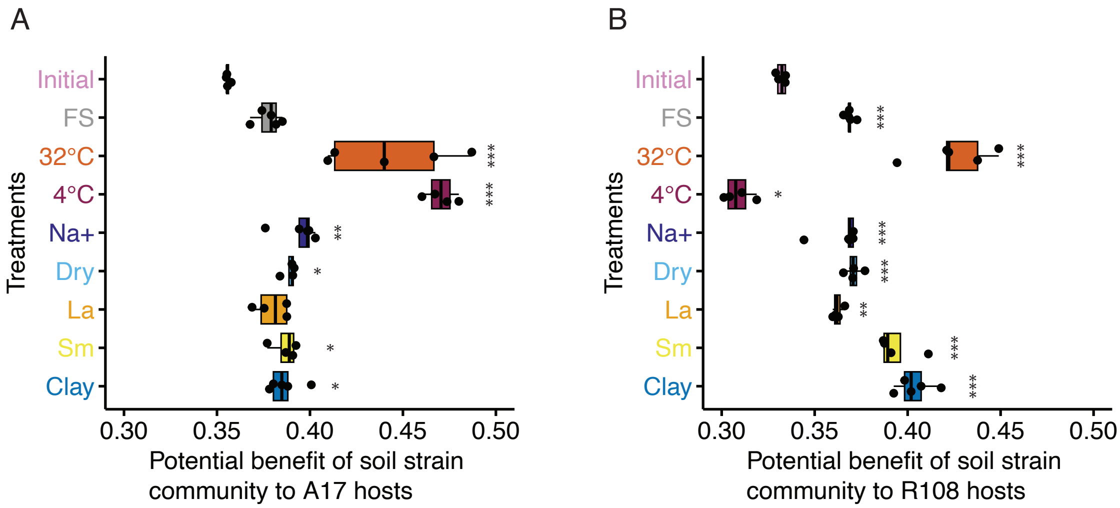
